# The Adaptor Protein Nck1, but not Nck2, Mediates Shear Stress-Induced Endothelial Permeability

**DOI:** 10.1101/651687

**Authors:** Mabruka Alfaidi, Umesh Bhattarai, Elizabeth D Cockerham, A.W. Orr

**Affiliations:** Department of Pathology and Translational Pathobiology, LSU Health – Shreveport, Shreveport, LA, USA; Department of Molecular & Cellular Physiology, LSU Health – Shreveport, Shreveport, LA, USA; Department of Cell Biology and Anatomy, LSU Health – Shreveport, Shreveport, LA, USA

**Keywords:** Shear stress, Vascular Permeability, Adaptor proteins Nck1 and Nck2, P-21 activated kinase

## Abstract

Alteration in hemodynamic shear stress at atheroprone sites promotes endothelial paracellular pore formation and permeability. Previously, we have reported that a peptide inhibitor to Nck prevented shear stress-induced p21 activated kinase (PAK) activation and endothelial permeability. However, the specificity of this peptide is unclear, and the role of individual Nck isoforms remain unknown. Here, we show that genetic deletion of Nck1/2 adaptor proteins significantly ameliorates shear stress induced permeability, and selective isoform depletion suggests distinct signaling mechanisms. Only Nck1 deletion significantly reduces flow-induced paracellular pore formation and permeability, whereas Nck2 depletion has no significant effects. Additionally, Nck1 reexpression, but not Nck2, restores shear stress-induced permeability in Nck1/2 knockout cells, confirming the non-compensating roles. *In vivo*, using the partial carotid ligation model of disturbed flow, Nck1 knockout prevented the increase in vascular permeability, as assessed by both Evans blue extravasation and leakage of plasma fibrinogen into the vessel wall. Domain swap experiments mixing SH2 (phosphotyrosine binding) and SH3 (proline rich binding) domains between Nck1 and Nck2 showed a dispensable role for SH2 domains but a critical role for the Nck1 SH3 domains in rescuing shear stress-induced endothelial permeability. Consistent with this, both Nck1 and Nck2 bind to PECAM-1 (SH2 dependent) in response to shear stress, but only Nck1 ablation interferes with shear stress-induced PAK2 activation (SH3 dependent). This work provides the first evidence that Nck1 and Nck2 play distinct roles in flow-induced vascular permeability.

**New and Noteworthy:** The present study shows a specific role for Nck1 in endothelial permeability in response to shear stress. Using *in vitro* and *in vivo* models, we demonstrate improvement in endothelial barrier integrity in cells subjected to disturbed flow only following Nck1 but not Nck2 deletion. Selective Nck1 inhibition may limit endothelial permeability at sites of disturbed flow to reduce atherosclerosis without affecting angiogenesis, which requires both Nck1 and Nck2 inhibition.

## Introduction

Shear stress regulates multiple aspects of atherosclerotic plaque formation through dynamic changes in endothelial permeability and leukocyte recruitment(25). Disturbed flow, observed at athero-prone regions, compromise endothelial layer integrity, resulting in enhanced permeability(4, 23). Multiple molecular mechanisms have been linked to flow-enhanced permeability, including activation of the serine/threonine kinase p21-activated kinase (PAK) in both *in vitro*(19) and *in vivo*(17) atherogenic models. Activated PAK localizes to cell-cell junctions and promotes endothelial permeability through cytoskeletal remodeling, turnover of junctional proteins, and paracellular pore formation(17, 24). However, the mechanisms by which PAK is recruited to cell-cell junctions in response to shear stress remains unknown.

The non-catalytic region of tyrosine kinase (Nck) family of adaptor proteins includes two isoforms (Nck1 and Nck2) composed exclusively of SH2/SH3 domains that lack enzymatic functions but mediate protein-protein interactions(1). Interactions between Nck’s SH2 domain and tyrosine phosphorylated proteins recruits Nck to sites of active cell signaling, where it brings in other signaling mediators through interactions between its SH3 domains and proline-rich sequences in target proteins(14). Nck signaling classically couples tyrosine phosphorylation to induction of actin cytoskeletal reorganization and cell movement(13). Consistent with this, genetic deletion of both Nck1 and Nck2, but not individual isoforms, impairs vascular development and endothelial angiogenesis(2). While the two isoforms of Nck were believed to have overlapping functions, non-compensating roles have recently been elucidated in multiple pathological settings. For example, Nck1 and Nck2 can act independently in dermal fibroblasts through distinct Rho GTPase family members; Nck1 promoted Cdc42 signaling for filopodium formation and Nck2 promoted Rho signaling to induce stress fibers(11).

Previous studies have shown that pretreating endothelial cells with a membrane-permeable peptide corresponding to the Nck-binding proline rich sequence of PAK (Pak-Nck peptide) reduces PAK targeting cell junction and blunts vascular permeability in atherosclerosis(5). However, the specificity of this peptide to Nck and the individual roles of Nck1 and Nck2 in shear stress-induced permeability remain unknown. Therefore, we utilized both cell culture and animal models of selective Nck1 and Nck2 depletion to characterize Nck signaling in flow-induced endothelial permeability.

## Materials and Methods

All reagents were provided from Gibco, USA, unless otherwise stated. All lentiviral vectors were designed and obtained using VectorBuilder website.

### Cell culture, Plasmids and RNA interference

Human aortic endothelial cells (HAECs, CELL Applications) were maintained in MCDB131 containing 10% (v/v) fetal bovine serum and cultured as we previously described(5). Nck1 and Nck2 were depleted from endothelial cells using lentiviral shRNA or CRISPR/Cas9 gene editing, or mouse endothelial cells were isolated from Nck1 or Nck2 deficient mice. The lentiviral vectors included pLV-(shRNA)-mCherry:T2A:Puro-U6 for Nck1 (target seq: GGGTTCTCTGTCAGAGAAA) and Nck2 (target seq: CTTAAAGCGTCAGGGAAGA) with 3^rd^ generation lenti components provided from Addgene; pMD2.G (12259), pRSV-Rev (12253), pMDLg/pRRE (12251). The lentiviral CRISPR/Cas9 plasmids utilized the pLV[CRISPR]-hCas9:T2A:Puro-U6>gRNA with the single-guide RNA (sgRNA) sequences targeting Nck1 (target seq: GTCGTCAATAACCTAAATAC), Nck2 (target seq: TGACGCGCGACCCCTTCACC) or scrambled sgRNA (target seq: GCACTACCAGAGCTAACTCA). The lentiviral vectors encoding full-length Nck1 and Nck2 were generated in the pLV-EXP-mCherry:T2A:Puro-CMV>Nck vector. For the domain swap experiments, pLV-EXP-mCherry:T2A:Puro-CMV>Nck constructs were generated containing the Nck1 SH2 domain and Nck2 SH3 domains or the Nck2 SH2 domain and Nck1 SH3 domains.

All cell lines were tested for the respective gene knocked down/knocked out before used for the experiments. Shear stress experiments were performed using the parallel plate flow chambers as we previously published(17, 18).

### Endothelial Permeability Assays

Endothelial cell permeability was determined by assessing paracellular pore formation following immunostaining for CD31 (abcam; ab9498, 5 µg/ml) or by using the Streptavidin/Biotinylated-Gelatin Trapping assay as previously described(9). Briefly, slides were coated with biotinylated Gelatin at 0.25 mg/ml overnight at 4°C. On the second day, endothelial cells were plated to confluence and subjected to oscillatory shear stress using a syringe pump (± 5dynes/cm^2^ with a superimposed 1dynes/cm^2^ for waste exchange for 18h). Immediately after cessation of flow, Streptavidin-Alexa Fluor 647 (1:1000, Invitrogen) was added to the cells for 1 minute before the addition of 3.7% (v/v) paraformaldehyde for the fixation. Alexa Fluor 488 Phalloidin (1:200, Invitrogen) was added to stain F-actin. For immunocytochemistry, cells were stained as previously described(5).

### Mouse experiments and in vivo permeability assay

All animal work was performed according to the National Research Council’s Guide for the Care and Use of Laboratory Animals and were approved by LSU Health-Shreveport Institutional Animal Care and Use Committee. Male ApoE^−/−^ mice on the C57BI/6J backgrounds were purchased from Jackson Laboratory (Bar Harvor, ME). Mice that contained alleles Nck1^−/−^ and Nck2^fl/fl^ were a gift from Tony Pawson (Lunenfeld-Tanenbaum Research Institute, Univ. of Toronto) whereas mice that contained Vascular Endothelial Cadherin (VE-Cad CreERT2 were kindly provided from Dr Luisa Iruela-Arispe, UCLA, CA). Mice were crossed with ApoE^−/−^ to generate endothelial specific (iEC) control mice (iEC-Control; ApoE^−/−^, VE-cadherin CreERT2^tg/?^), Nck1 knockout (KO) mice (ApoE^−/−^, VE-cadherin CreERT2^tg/?^, Nck1^−/−^), endothelial-specific Nck2 knockout mice (iEC-Nck2 KO; ApoE^−/−^, VE-cadherin CreERT2^tg/?^, Nck2^fl/fl^), and endothelial-specific Nck1/2 double knockout (DKO) mice (iEC-Nck1/2 DKO; ApoE^−/−^, VE-cadherin CreERT2^tg/?^, Nck2^fl/fl^, Nck1^−/−^). At 8-9 weeks of age, mice were intraperitoneally injected with Tamoxifen (1 mg/kg, Sigma, St Louis, MO) for five subsequent days to induce Cre expression and gene excision. After 2 week recovery, the animals were subjected to partial carotid ligation surgery as previously described(15). One week post-ligation, mice were injected with 1% (w/v) Evans Blue (Sigma, E2129) retro-orbitally. After 30 minutes, the animals were euthanized and left and right carotid arteries were collected. Evans blue extravasation in left and right carotid arteries were assessed as previously described(22).

### Immunohistochemistry

Sections of carotid arteries were stained as previously described(26). Heat mediated antigen retrieval pretreatment using 10 mM Sodium Citrate buffer (Vectors Biolabs) were used. Primary antibodies (CD31, sc-1506, 1 µg/ml, Fibrinogen, abcam, ab34269, 5 µg/ml) were incubated at 4°C overnight. Images were captured with a Nikon microscope and analyzed using Nis Elements software. The investigators were blinded to the animal groups during the process of data collection and analysis.

### Statistical analysis

Data are analyzed using GraphPad Prism as mean ± SEM. Data was first tested for Normality (Kolmogorov-Smirnov test). For multiple comparison, one or Two Way ANOVA, followed by Bonferroni’s post-test. Statistical significance was achieved only when p<0.05.

## Results

### Nck1/2 are required for shear stress induced permeability

To test whether Nck1 and Nck2 are involved in shear stress-induced permeability, we first deleted Nck1 and Nck2 genes in endothelial cells using the CRISPR/Cas9 system by sequential deletion of Nck1 and Nck2 (Nck1/2 DKO). Complete endothelial Nck1 and Nck2 deletion was confirmed using Western blot (Figure 1A). We previously demonstrated that shear stress increased endothelial permeability by paracellular pore formation(19). Endothelial cells subjected to acute shear stress for 30 minutes showed a significant increase in paracellular pore formation that was absent in the Nck1/2 DKO cells (Figure 1B-C). Endothelial permeability was also assessed by 647 Alexa-streptavidin-biotinylated gelatin trapping assay. While there were no significant differences between the two cell types (scrambled gRNA vs. Nck1/2 DKO) at basal levels, there was remarkable reduction in the shear-induced permeability in Nck1/2 DKO cells following chronic oscillatory flow (Figure 1D/E), suggesting that Nck1/2 adaptor proteins critically couple shear stress to endothelial permeability.

**Figure 1.**
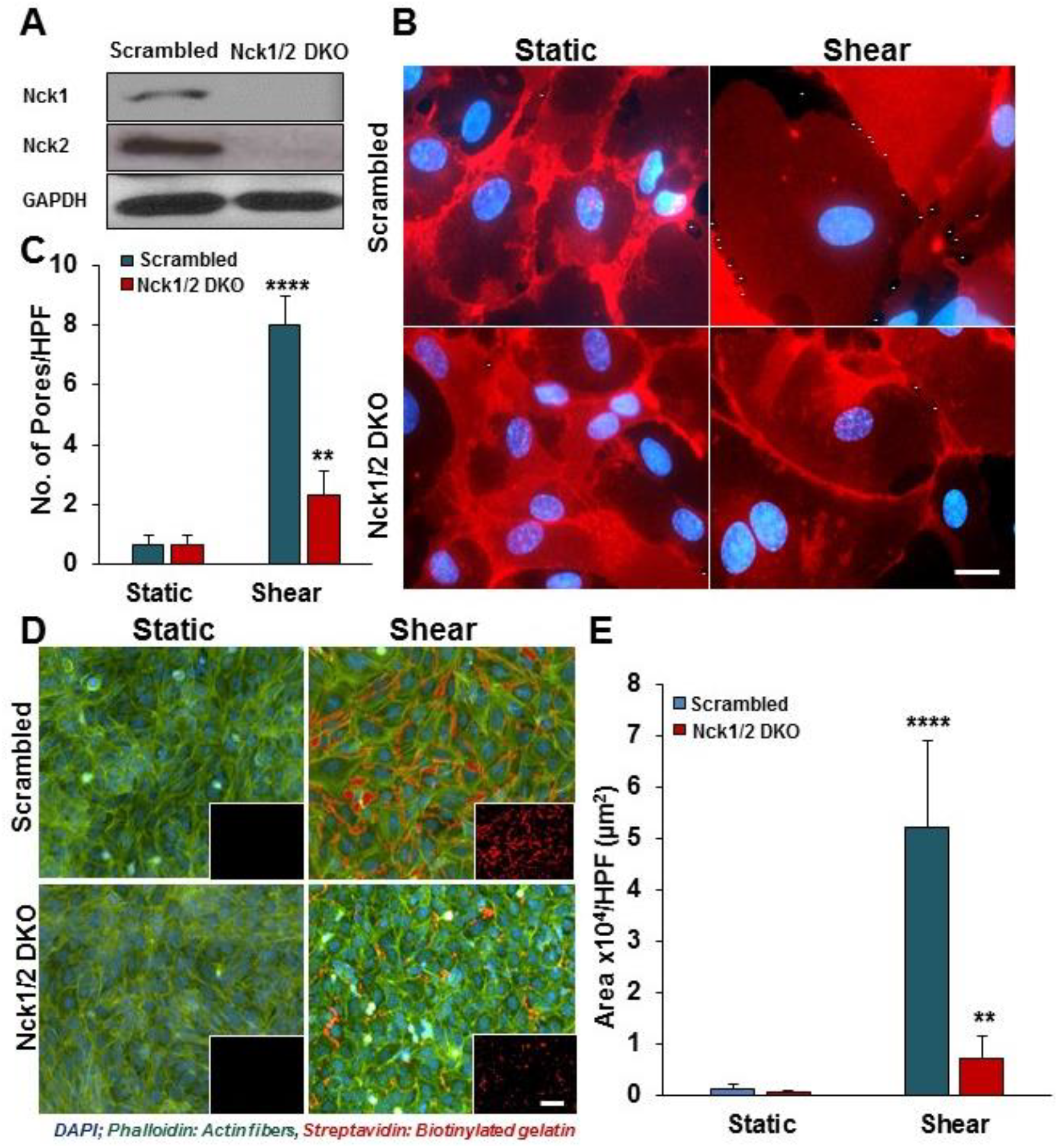
Deletion of Nck1 and Nck2 adaptor proteins blunt shear stress-induced endothelial permeability. **A)** A representative blot showing deletion of Nck1 and Nck2 in Nck1/2 DKO cells. **B-C)** endothelial monolayer integrity was assessed in Nck1/2 DKO or Scramble cells after shear stress (30 minutes) and stained with PECAM-1 (Red) for paracellular pore formation (stars). **D-E)** Endothelial cell permeability assessed after oscillatory shear exposure (OSS; ±5 dynes/cm2 with 1dynes/cm2 superimposed flow) for 18h, using the biotinylated gelatin-Alexa647 Streptavidin (Red) system, F-actin was labeled with Alexa488-Phalloidin (Green). All data are from n=4, mean ± SEM, analyzed by 2-way ANOVA, and Bonferroni’s post-test, **p<0.01. Scale bars=100-200µm.

### Nck1 but not Nck2 ablation decreases shear stress-induced permeability

Having shown that deletion of both Nck1 and Nck2 decreases shear stress induced permeability, we sought to determine whether individual Nck isoforms differentially mediate this response. Using lentiviral shRNA delivery, we specifically knocked down Nck1 and Nck2 in endothelial cells and confirmed the depletion was isoform-specific (Figure 2A). Only Nck1 deleted cells showed significant amelioration of shear stress-induced paracellular pore formation, whereas Nck2 ablated cells showed enhanced pore formation (Figure 2B-C). Similarly, the shear-induced increase in streptavidin leak was significantly decreased in Nck1, but not Nck2, depleted cells (Figure 2D). Acute siRNA-mediated Nck1 and Nck2 knockdown produced similar results (data not shown), confirming that the Nck1 isoform rather than Nck2 isoform regulates endothelial permeability by flow.

**Figure 2.**
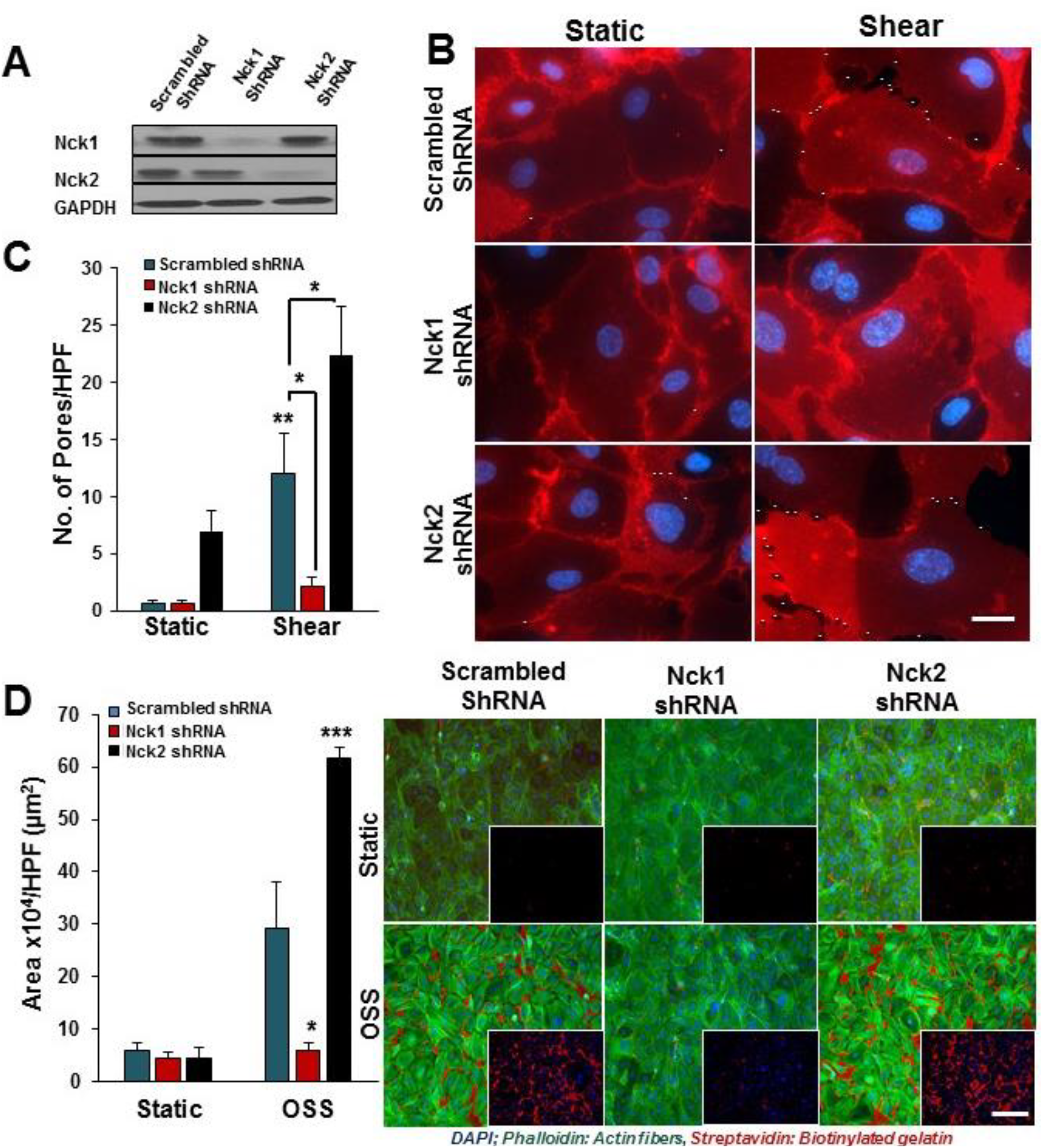
Ablation of Nck1 but not Nck2 adaptor protein blunts shear stress-induced endothelial permeability. **A)** Representative blots showing the efficiency of the knocking down in HAECs after Nck1 shRNA, Nck2 shRNA, or scramble control transduction. (**B &C)** PECAM-1 stained cells showing paracellualr formation (arrows). **D)** Nck1 but not Nck2 knocking down prevents flow induced permeability using biotinylated gelatin/Alexa647-Streptavidin assay. DAPI (blue; Nuclei), Phalloidin (Green; F-actin Fibers), Streptavidin (Red; Biotinylated Gelatin). Images analyzed using Nis Elements software from n=4. Scale bars=100-200µm. Data are analyzed by 2-Way ANOVA and Bonferroni’s post-test, *p<0.05.

*In vivo*, areas of disturbed flow show elevated endothelial permeability(12). Therefore, to study the effects of Nck1 and Nck2 on shear stress-induced permeability *in vivo*, we elected to perform partial carotid ligation (PCL) to induce disturbed flow in the left common carotid as previously described(REF). iEC-Control, Nck1 KO, iEC-Nck2 KO, and iEC-Nck1/2 DKO mice were subjected to PCL and the left carotid was exposed to disturbed flow for 7 days. The unligated right carotid serves as an internal control for non-disturbed flow. Disturbed blood flow after ligation was confirmed using echocardiography (data not shown). After 1 week, permeability was assessed by Evans Blue Dye injection (Figure 3A). While there were no significant changes in Evans blue extravasation in the right (un-ligated) carotids among all experimental groups, only Nck1 KO and iEC-Nck1/2 DKO mice showed a two-fold reduction in Evans blue accumulation in the ligated left carotid (Figure 3B). To verify this isoform selective permeability effect, permeability was also assessed by measuring leak of the plasma protein fibrinogen (FG) into the ligated carotid arteries. Consistent with previous data, the leakage of plasma FG was significantly ameliorated in Nck1 KO and iEC-Nck1/2 DKO mice compared to controls. However, Nck2 KO mice did not significantly affect either Evans Blue extravasation or FG staining (Figure 3C-D).

**Figure 3.**
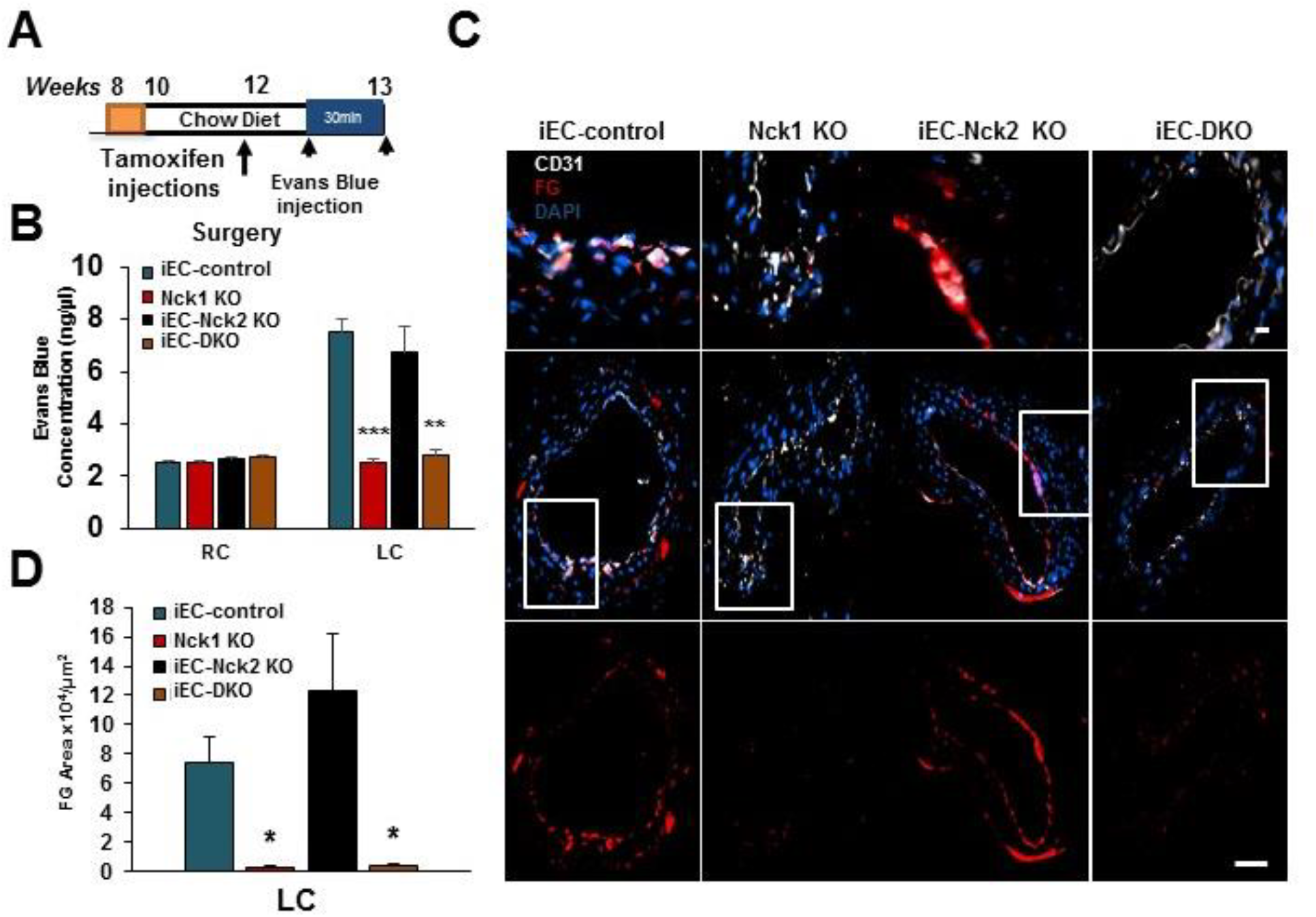
Ablation of Nck1 but not Nck2 adaptor protein blunts Partial Carotid ligation induced permeability. **A)** Schematic of the study in which four groups of mice were subjected to the ligation surgery as indicated animal genotypes and time of surgery. **B)** Nck1 but not Nck2 knockout mice showed less Evans Bule extravasation and **(C-D)** fibrinogen (FG: Red) staining in ligated left carotid arteries. Images analyzed using Nis Elements software, from n=7-10 mice/group. Data are mean ± SEM, analyzed by1-way or 2-way ANOVA, and Bonferroni’s post-test, *p<0.05, **p<0.01, ***p<0.001

### Nck1 regulates shear stress-induced permeability by its SH3 domains

To characterize why Nck1 but not the highly homologous Nck2 is capable of regulating endothelial barrier function in response to shear stress, we conducted domain swap experiments with chimeric proteins in which the Nck1 SH2 domain was paired with Nck2 SH3 domains or the Nck2 SH2 domain was paired with the Nck1 SH3 domains (Figure 4A). The expression of full length Nck1, Nck2, and the chimeric proteins Nck1 SH2/Nck2 SH3 and Nck2 SH2/Nck1 SH3 were confirmed by mCherry fluorescence (data not shown) and Western blotting (Figure 4B). As expected, the re-expression of Nck1 but not Nck2 in Nck1/2 DKO cells rescued the permeability response to oscillatory shear stress (Figure 4C/D). The chimera containing the Nck1 SH2 domain and Nck2 SH3 domains failed to rescue the permeability response, suggesting that Nck1 SH3 domains are critical to the permeability response (Figure 4 C/D). The chimera containing the Nck2 SH2 and Nck1 SH3 domains significantly enhanced flow-induced endothelial permeability, suggesting that the Nck1 and Nck2 SH2 domains are both capable of targeting Nck to the proper location for SH3 domain-mediated signaling (Figure 4C-D). Together, these data reveal that Nck1 SH3 domains (1-3) mediate the isoform-selective effects of Nck1 on shear stress-induced permeability, whereas the SH2 domains of Nck1 and Nck2 are redundant.

**Figure 4.**
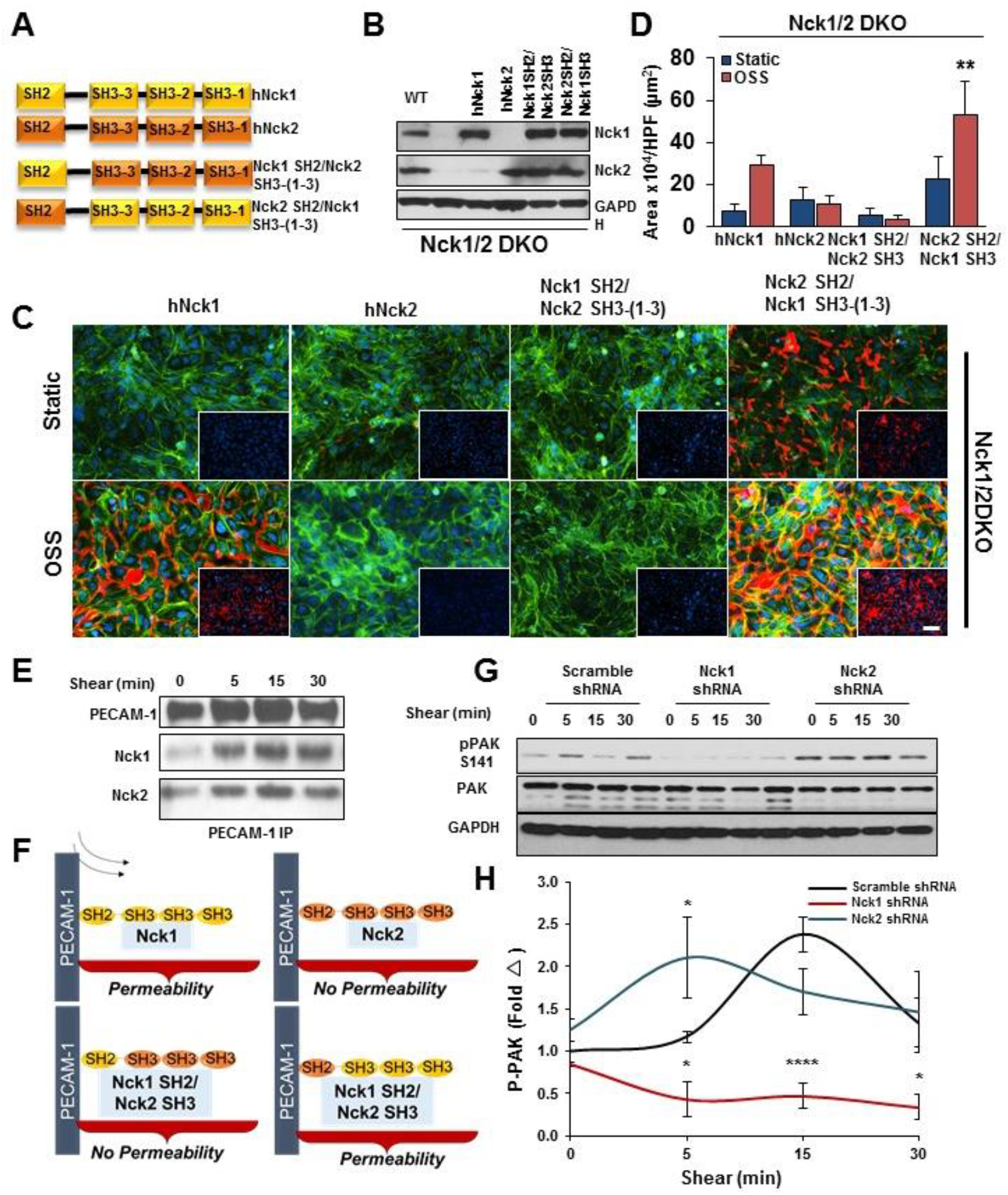
SH3 domains of Nck1 are essential for flow induced endothelial permeability. **A)** Schematic showing the domain structure of Nck1 and Nck2 and two chimera of Nck1 SH2/ SH3, and Nck1 SH3/ Nck2 SH2. **B)** Western blot analysis showing comparable Nck1/2 expression following transduction of constructs in **(A)** in Nck1/2 DKO cell lysates. **C-D)** permeability of endothelial cells after oscillatory shear stress (OSS) as assessed by Streptavidin-Biotinylated Gelatin trapping assay. Data are analyzed by 2-Way ANOVA, and Bonferroni’s post, n=4, **p<0.01. **E)** Nck1 and Nck2 in PECAM-1 co-immunoprecipitation assay confirms the direct interactions between Nck1/2 with PECAM-1 in response to shear stress. From n=4 independent experiments. **F)** Schematic diagram suggesting that Nck1/2 bind to PECAM-1 in response to shear stress, but only Nck1 through its SH3 domains regulate the permeability. **G-H)** PAK2 activation as assessed by phosphorylation is downregulated by Nck1 ablation, data are mean ± SEM, analyzed by One-Way ANOVA, *p<0.05, ****p<0.0001.

### Nck1 mediates permeability by promoting PAK recruitment to PECAM-1

We previously showed that tyrosine phosphorylated platelet endothelial cell adhesion molecule-1 (PECAM-1) in the endothelial adherens junction could recruit Nck1/2 in response to oxidative stress(3). Co-immunoprecipitation assays confirmed that both Nck1 and Nck2 interacted with PECAM-1 following shear stress in a time dependent manner (Figure 4E). These data are consistent with redundant SH2 domains recruiting both Nck1 and Nck2 to PECAM-1, whereas the Nck1 SH3 domains are required to induce endothelial permeability (Figure 4F). Since previous data pointed to a role for PAK in shear stress induced endothelial permeability(19) and PAK binds to the second SH3 domain of Nck(10), we elected to assess PAK activation in response to shear stress. Shear stress induced PAK activation (measured by Ser141 phosphorylation) was significantly diminished in Nck1 ablated cells (Figure 4G-H). By contrast, Nck2 depleted cells showed enhanced basal PAK phosphorylation consistent with enhanced endothelial permeability (Figure 4G-H). Taken together, the data suggest that Nck1 selectively mediates flow-induced PAK activation and induction of endothelial permeability.

## Discussion

Endothelial cell barrier function is important to maintain the integrity of blood compartment and permit passage of soluble factors in a tightly regulated manner(27). Hemodynamic shear stress acts locally and regulates vascular permeability by controling cytoskeletal remodeling and endothelial cell-cell junction stability(20). While laminar flow improves the barrier integrity, dysrgulation of this barrier has been obsreved at atheroprone areas secondary to disturbed oscillatory blood flow(7). Multiple pathways have been implicated, however, the underlying molecular mechanims remain largely unknown. In the present study we have shown that formation of paracellular pores and subsequently enhanced permeability in response to shear stress is prevented by selective depletion of Nck1. Mechanistically, both Nck1 and Nck2 bind to PECAM-1 but the differential Nck1 effects on the barrier function was mediated by the SH3 domains and isoform-selective PAK activation. We and others have previously reported that endothelial cells treated with a cell permeable peptide containing the Nck-binding sequence from PAK inhibits flow-induced permeability *in vitro*(24) and *in vivo*(19). However, these studies couldn’t determine whether this peptide acts specifically through blocking the PAK-Nck interaction. Herein, we showed for the first time that Nck adaptor proteins are critical for flow-induced endothelial permeability and para-cellular pore formation.

The two highly similar Nck proteins (Nck1 and Nck2) are expressed by different genes(21) that play redundant roles during development, as deletion of both Nck isoforms results in an embryonic lethal phenotype due to impaired vasculogenesis while deletion of only one isoform did not(2). While Nck1 and Nck2 play redundant roles in regulating angiogenesis in mouse models of retinopathy(8), Nck2 may play a dominant role in PDGF-induced actin polymerization in NTH3T3(6) and nerve growth factor-induced axon and dendrite tree in primary rat cortical neurons(11). In contrast, Nck1 plays a more dominant role in T cell receptor-induced ERK activation and inflammation(16), suggesting non-compensating roles during phenotypic regulation post-development and points to differences in specific binding affinities between Nck1 and Nck2, despite their overall sequence homology(13). We now demonstrate a non-redundant role for Nck1 in endothelial permeability by showing that despite Nck1/2 binding to PECAM-1 only Nck1 SH3 (1–3) are involved in regulating endothelial permeability. The critical role for Nck1 SH3 domains appears to involve selective regulation of flow-induced PAK activation by Nck1. However, both Nck1 and Nck2 have been shown to interact with PAK, and the mechanisms by which Nck1 critically regulates PAK activation by flow remains unknown.

Multiple studies have shown that treatment with a PAK-Nck blocking peptide reduces permeability and inflammation in a variety of pathological conditions(5, 19). However, Nck adaptor proteins critically regulate angiogenesis, suggesting that Nck inhibition would be detrimental in conditions where angiogenesis promotes proper healing, such as ischemic injury. Since multiple groups have shown that both Nck1 and Nck2 must be inhibited to reduce angiogenesis(8), these data suggest that selective inhibition of Nck1 may therefore allow for improved barrier function without compromising the angiogenic response.

## Acknowledgments

The authors thank Dr. Tony Pawson (Lunenfeld-Tanenbaum Research Institute, Univ. of Toronto) (Nck1ko and Nck2^flox/flox^ mice), and Dr. Luisa Iruela-Arispe (UCLA, CA) (VeCad Cre mice).

